# An influenza A hemagglutinin small-molecule fusion inhibitor identified by a new high-throughput fluorescence polarization screen

**DOI:** 10.1101/2020.04.02.022160

**Authors:** Yao Yao, Rameshwar U. Kadam, Chang-Chun David Lee, Jordan L. Woehl, Nicholas C. Wu, Xueyong Zhu, Seiya Kitamura, Ian A. Wilson, Dennis W. Wolan

## Abstract

Influenza hemagglutinin (HA) glycoprotein is the primary surface antigen targeted by the host immune response and a focus for development of novel vaccines, broadly neutralizing antibodies (bnAbs) and therapeutics. HA enables viral entry into host cells via receptor binding and membrane fusion and is a validated target for drug discovery. However, to date, only a very few bona fide small molecules have been reported against the HA. To identity new antiviral lead candidates against the highly conserved fusion machinery in the HA stem, we synthesized a fluorescence-polarization probe based on a recently described neutralizing cyclic peptide P7 derived from the complementarity-determining region loops of human bnAbs FI6v3 and CR9114 against the HA stem. We then designed a robust binding assay compatible with high-throughput screening to identify molecules with low μM to nM affinity to influenza A group 1 HAs. Our simple, low-cost, and efficient in vitro assay was used to screen H1/Puerto Rico/8/1934 HA trimer against approximately 72,000 compounds. The crystal structure of H1/Puerto Rico/8/1934 HA in complex with our best hit compound F0045(S) confirmed that it binds to pockets in the HA stem similar to bnAbs FI6v3 and CR9114, cyclic peptide P7, and small molecule inhibitor JNJ4796. F0045 is enantioselective against a panel of group 1 HAs and F0045(S) exhibits in vitro neutralization activity against multiple H1N1 and H5N1 strains. Our assay, compound characterization, and small-molecule candidate should further stimulate the discovery and development of new compounds with unique chemical scaffolds and enhanced influenza antiviral capabilities.

**Summary:** Influenza hemagglutinin (HA) glycoprotein enables viral entry into host cells and is the main target for antibodies in our immune system. While HA has now been established as a validated target for drug discovery, no FDA-approved small molecules are available that specifically prevent HA from binding host receptors or inhibit its membrane fusion activity and thus prevent infection. We therefore designed a fluorescence polarization probe to enable rapid identification of small molecules that bind to the stem fusion machinery of group 1 HAs. Application of our assay yielded a small molecule to the influenza A group 1 HA stem with antiviral efficacy.

## Introduction

Influenza, together with subsequent complications by bacterial pneumonia (*i.e., Streptococcus pneumoniae, Staphylococcus aureus*) is among the top 10 leading causes of death in the US with over 50,000 people succumbing to infection each year. According to Centers for Disease Control and Prevention, the estimated number of U.S. hospitalizations and deaths directly due to influenza during the 2017-2018 season was 960,000 and 79,000, respectively (1). More devasting are the unpredictable pandemic strains that can result in the mortality of one million (1957 Asian flu) to over 50 million (1918 Spanish flu) individuals (2). Thus, there is a critical need for readily available therapeutics to combat the global spread of influenza. Currently, three types of FDA-approved anti-influenza drugs are available and include: i) neuraminidase inhibitors, such as oseltamivir (Tamiflu) and zanamivir (Relenza), that prevent release of nascent virions post-infection (3); ii) M2 ion channel inhibitors amantadine (Symmetrel) and rimantadine (Flumadine) that act by preventing viral uncoating during early stages of replication (4); and iii) the cap-dependent endonuclease inhibitor baloxavir marboxil (Xofluza), which is the most recently US FDA approved drug for influenza A and B viruses (5). Unfortunately, all of these molecules are subject to rapid resistance by influenza viruses and resistant clinical isolates have been reported (6-8). Thus, there is a pressing and unmet need for new broad-spectrum influenza antivirals to combat pandemics and seasonal epidemics that can work alone or synergistically with current therapeutics and/or the host adaptive immune system.

The hemagglutinin (HA) glycoprotein is the most abundant transmembrane protein on the surface of influenza and is necessary for initiating viral infection. HA is a trimeric class I viral fusion protein with a globular membrane-distal head containing the receptor binding site (RBS) and a membrane-proximal stem housing the fusion machinery (9). These two regions of HA have two essential functions of facilitating entry into host cells through binding of HA to host sialosides (10) and membrane fusion upon entry into endosomes (11), respectively. Endosomal uptake is necessary for infection, where the low pH of the endosome triggers conformational rearrangements in the metastable prefusion HA that lead to a post-fusion state and subsequent membrane fusion (12, 13). Importantly, a number of broadly neutralizing antibodies (bnAbs) have now been characterized, including CR9114 and FI6v3 (14, 15), that bind and stabilize the HA stem and prevent these conformational rearrangements in low pH conditions. Stabilization of the pre-fusion state HA by small molecules would then mimic the stem bnAbs and impede membrane fusion to effectively prevent infection (16).

Despite HA now being an established drug target, no FDA-approved therapeutics specifically block receptor binding or the fusion machinery. Umifenovir (Arbidol) is an antiviral small molecule used to treat influenza and other respiratory diseases in Russia and China (17, 18); however, large doses are required to achieve therapeutic efficacy (19). While Umifenovir also has broad-spectrum antiviral capabilities against Ebola, hepatitis B, and hepatitis C (17, 18), we recently demonstrated that it binds to a specific hydrophobic cavity in the upper region of the HA stem and this binding site likely accounts for its influenza antiviral effects (20). Notwithstanding, Umifenovir needs extensive optimization to improve HA affinity and pharmacokinetic stability and additional molecules that target HA at other surface epitopes are urgently needed.

We and colleagues recently reported a small cyclic peptide P7 that neutralizes group 1 HA influenza (21). P7 is based on the heavy-chain complementarity-determining region (HCDR) loop of bnAb FI6v3 and framework region 3 (FR3) of CR9114. Similar to bnAb FI6v3, crystal structures and cellular assays demonstrate P7 binds to the highly conserved HA stem epitope and blocks the low pH–induced conformational rearrangements associated with membrane fusion. However, unlike stem-targeted bnAbs FI6v3 and CR9114, P7 is specific to influenza A group 1 HAs. The P7 specificity for group 1 is due in part to glycosylation of Asn38 in group 2 HAs as well as substitution of Thr49 (group 1) for a larger Asn (group 2) that interferes with binding. In another collaboration with Janssen, we also disclosed the small molecule JNJ4796, an HA stem-targeted inhibitor that is orally active in mice (22). JNJ4796 can neutralize a broad spectrum of human pandemic, seasonal, and emerging group 1 influenza A viruses and has promise as a therapeutic option with a complementary mechanism-of-action to existing antiviral drugs for the treatment of influenza.

The primary issue with the identification of HA-targeted small molecules is the lack of robust, rapid, and cost-effective high-throughput screening (HTS) assays (22-24). Identification of new antiviral small molecules have been attempted with both phenotypic and ELISA assay formats, including a dual myxovirus reporter assay (23) and an AlphaLISA assay (amplified luminescent proximity homogeneous assay) (22). The AlphaLISA screen was used to identified the precursor hit molecule of JNJ4796, whereby a diverse library of ∼500,000 compounds was screened for displacement of the *de novo*-designed small protein HB80.4 (25) from the group 1 HA stem epitope. While these assays helped to identify new lead antiviral candidates, AlphaLISA and phenotypic HTS formats can be expensive, convoluted, and difficult to develop counter-screens to eliminate false positives.

We present here the design, synthesis, and application of a P7 peptide-based fluorescence polarization (FP) probe that is selective for the screening of the stem epitope of influenza A group 1 HAs. As a proof-of-concept, we performed an HTS against 72,000 compounds to identify molecules with affinity to the H1/Puerto Rico/8/1934 (H1/PR8) HA stem. We identified a novel small molecule F0045(S) and biophysically characterized its binding to a panel of group 1 HAs with surface plasmon resonance, x-ray crystallography, and differential scanning fluorimetry. Importantly, F0045(S) neutralizes influenza A infection and our assay and hit molecule thus represent key advancements in the ability to interrogate HA for additional molecular scaffolds that can be optimized for development of antiviral influenza drugs.

## Results and Discussion

Fluorescence polarization is a powerful approach by which alterations in the apparent molecular weight of a fluorescent probe in solution are indicated by changes in the polarization of emitted light from the sample (26). A robust FP probe should ideally result in low and high FP when incubated alone or in the presence of a protein target, respectively (**Fig. 1A**). Our FP probe was designed based on functional and structural considerations of the cyclic peptide P7 that targets the highly conserved stem epitope of influenza A group 1 HA with low nanomolar binding affinity. Based on the crystal structure of H1 HA in complex with P7 (21), we attached a carboxytetramethylrhodamine (TAMRA) fluorophore to a free amine located on the P7 aminopropanamide moiety, as this side chain is located in a solvent channel with limited interactions to HA. We posited that modification of the cyclic peptide at this position would have minimal interference on P7 binding to HA (**Fig. 1B**). The P7-TAMRA probe was synthesized by a direct amine-carboxylate acid coupling between the pure P7 peptide (synthesized by standard Fmoc-based solid-phase peptide synthesis procedures) and TAMRA. The P7-TAMRA probe was purified by reverse-phase HPLC purification to yield >95% product (see Materials and Methods for additional details).

**Figure 1.**
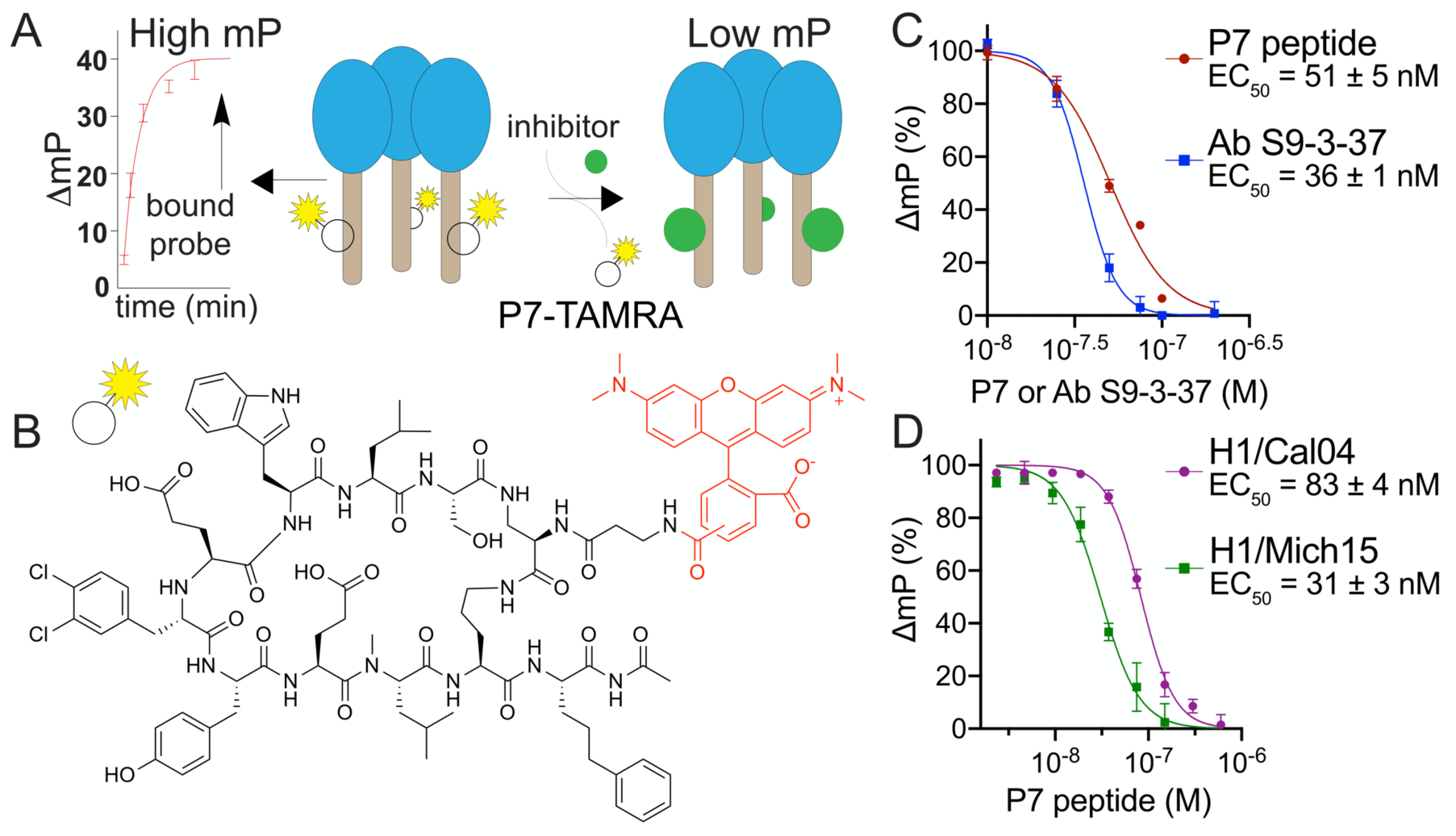
Design and characterization of the P7-based FP probe. (A) Design of FP assay to identify molecules with affinity to the HA stem. The polarization signal of the P7-TAMRA probe produces high and low FP when bound or unbound to the HA protein, respectively. (B) Chemical structure of P7-TAMRA probe. (C) P7-TAMRA probe (75 nM) and H1/PR8 (30 nM) competition assay against a dose-dependent increase in P7 peptide or stem-targeting bnAb S9-3-37 (mP, millipolarization). (D) P7-TAMRA probe (75 nM) and other HA (H1/Cal04 (30 nM) or H1/Mich15 (100 nM)) competition assays against P7 peptide.

We first optimized the P7-TAMRA probe and H1/PR8 HA concentrations to yield the best separation between high and low FP signals in a 96-well plate format in the presence and absence of protein. Importantly, fluorescence polarization of the P7-TAMRA probe increased only in the presence of H1/PR8 HA, while no observable change in polarization was detected when the P7-TAMRA probe was incubated with the group 2 A/Hong Kong/1/1968 (H3/HK68) HA (**Fig. S1**). We next assessed the ability of P7 to compete with the P7-TAMRA probe for HA stem binding. Increasing concentrations of P7 were introduced to pre-incubated solutions of 30 nM H1/PR8 HA and 75 nM P7-TAMRA in a buffer consisting of PBS, pH 7.4 and 0.01% Triton X-100 in a final volume of 60 μL. After a 1-min incubation at room temperature, fluorescence polarization was measured on a PerkinElmer EnVision plate reader and we calculated a relative EC_50_ of 51 ± 5 nM (**Fig. 1C**). This value is comparable to that for P7 (EC_50_ = 30 to 70 nM) that was previously measured by an AlphaLISA assay using the small protein HB80.4 as a competitor for HA binding (21). In addition, we also performed the competition assay with a group 1-specific bnAb S9-3-37 (27) and measured an EC_50_ of 36 ± 1 nM (**Fig. 1C**). The FP assay was next assessed for adaptability and sensitivity to measure FP of the P7-TAMRA probe in the presence of other group 1 HAs, including A/California/04/2009 (H1/Cal04) and A/Michigan/45/2015 (H1/Mich15). A repeat of the P7 vs. P7-TAMRA competition assay with these HAs yielded relative EC_50_ values of 83 ± 4 nM and 31 ± 3 nM, respectively (**Fig. 1D**).

We next sought to optimize our cost-effective and throughput assay to identify new small molecules with affinity to the HA stem epitope. Similar to the proof-of-concept FP assays that used 30 nM H1/PR8 HA and 75 nM P7-TAMRA probe, we miniaturized the volume to 10 μL to perform high-throughput screening (HTS) against commercially available small molecule libraries. The assay was optimized for maximum differential FP between negative and positive controls consisting of DMSO and 300 nM P7 peptide, respectively (**Fig. 2A**). A simplified two-step HTS process was performed, whereby 100 nL of 2 mM DMSO stock solutions of small molecules was introduced into a pre-mixture of HA and P7-TAMRA distributed into 384-well low-volume plates (final compound concentration was 20 μM). Plates were incubated for 30 min at room temperature and subsequently analyzed for FP, as described in Materials and Methods. We screened H1/PR8 HA against 72,000 compounds consisting of the commercially available Maybridge HitFinder, ChemDiv, and Life Chemical libraries (**Fig. 2A**). The average Z’ for the HTS was 0.81 with an overall hit rate of 0.01%, where molecules with FP < 3x the coefficient of variation (CV) were considered hits (28, 29) (**Fig. 2A**). Importantly, our HTS assay has significant advantages over previous screening attempts, as the FP can be rapidly measured upon reagent mixing, cost-effectiveness due to the limited amount of reagents required (*i.e.*, 7.5 nmol P7-TAMRA probe and 3 nmol HA per 10,000 compounds), and ease of removing false-positive hits that are typically due to inherent fluorescence.

**Figure 2.**
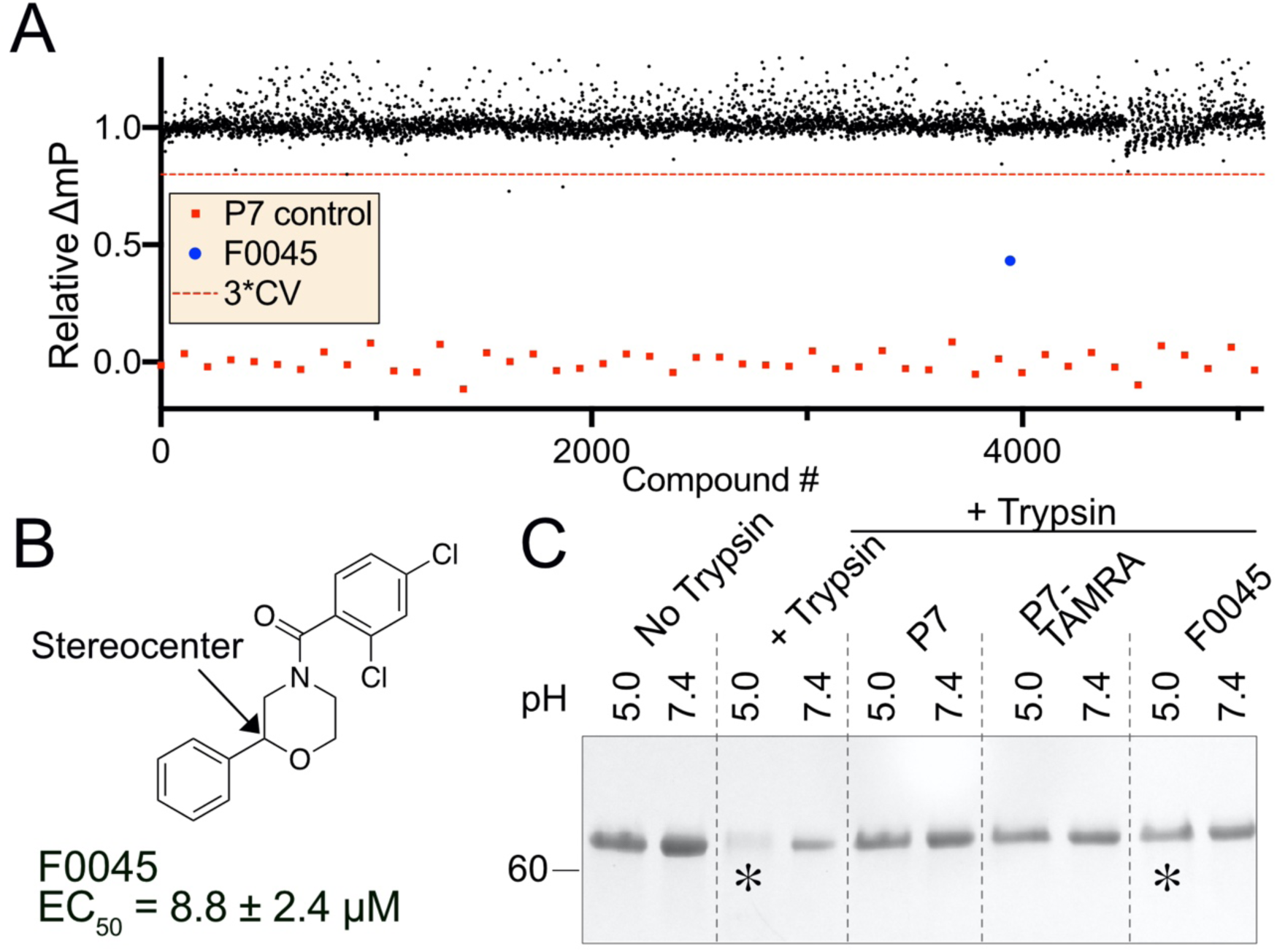
HTS for H1/PR8 and hit confirmation. (A) Scatter plot representation of 15 384-well plates in HTS with F0045 shown in blue. Black dots represent number of compounds screened and red dots as a peptide P7 control (CV, coefficient of variation). (B) Chemical structure of the best hit F0045 from the HTS campaign. (C) SDS/PAGE analysis of the trypsin-susceptibility assay of 5 μM H1/PR8 HA incubated with 50 μM P7, P7-TAMRA probe, or F0045 with DMSO as a negative control.

All library wells consisting of compounds with significantly reduced FP were submitted for LC-MS/MS verification and molecules were purchased from vendors for further *in vitro* characterization. One small molecule, termed F0045, was subjected to 8-point dose-response assays with H1/PR8 HA and the resulting EC_50_ of 8.8 ± 2.4 μM provided confidence that this molecule was an authentic hit worth further analysis (**Figs. 2B, S2**). We next employed an established trypsin susceptibility assay to assess if F0045 protected H1/PR8 from low pH induced conformational changes, akin to the endosomal membrane fusion event (20-22). Briefly, H1/PR8 HA is susceptible to trypsin digestion in the low pH post-fusion state, but stem-binding small molecules (*i.e.*, Umifenovir, JNJ4796) or bnAbs (*i.e.*, FI6v3, CR9114) prevent the HA from undergoing these conformation rearrangements on lowering of the environmental pH (**Fig. 2C**). Similar to the P7 peptide and P7-TAMRA probe, 50 μM F0045 prevented 5 µM H1/PR8 HA from trypsin digestion suggesting that the small molecule prevents HA from transitioning to the post-fusion state at pH 5.0. Thus, F0045 has reasonable *in vitro* HA affinity in the low μM range and effectively prevents the biologically relevant low-pH HA conformational change.

Our best hit F0045 has a stereocenter and the compound within the commercial library likely consisted of a racemic mixture (**Fig. 2B**). We further investigated if the spatial orientation of the phenyl group has a role in HA affinity by synthesizing both F0045 enantiomers (**Fig. 3A**, see Supplementary Information for synthesis). The S-enantiomer (*e.g.*, F0045(S), EC_50_ = 1.9 ± 0.3 μM) has a significantly reduced relative EC_50_ than the R-enantiomer (*e.g.*, F0045(R), EC_50_ = 43 ± 8 μM) when measured by our FP competition assay using the P7-TAMRA probe and H1/PR8 HA (**Fig. 3B**). A similar trend of improved affinity by the S-enantiomer F0045(S) was also observed for the HAs of other H1 strains, including H1/Cal04 (EC_50_ = 1.4 ± 0.9 μM) and H1/Mich15 (EC_50_ = 0.50 ± 0.16 μM). The spatial orientation of the phenyl group then has a significant effect on the compound’s ability to bind HA (**Fig. 3B**).

**Figure 3.**
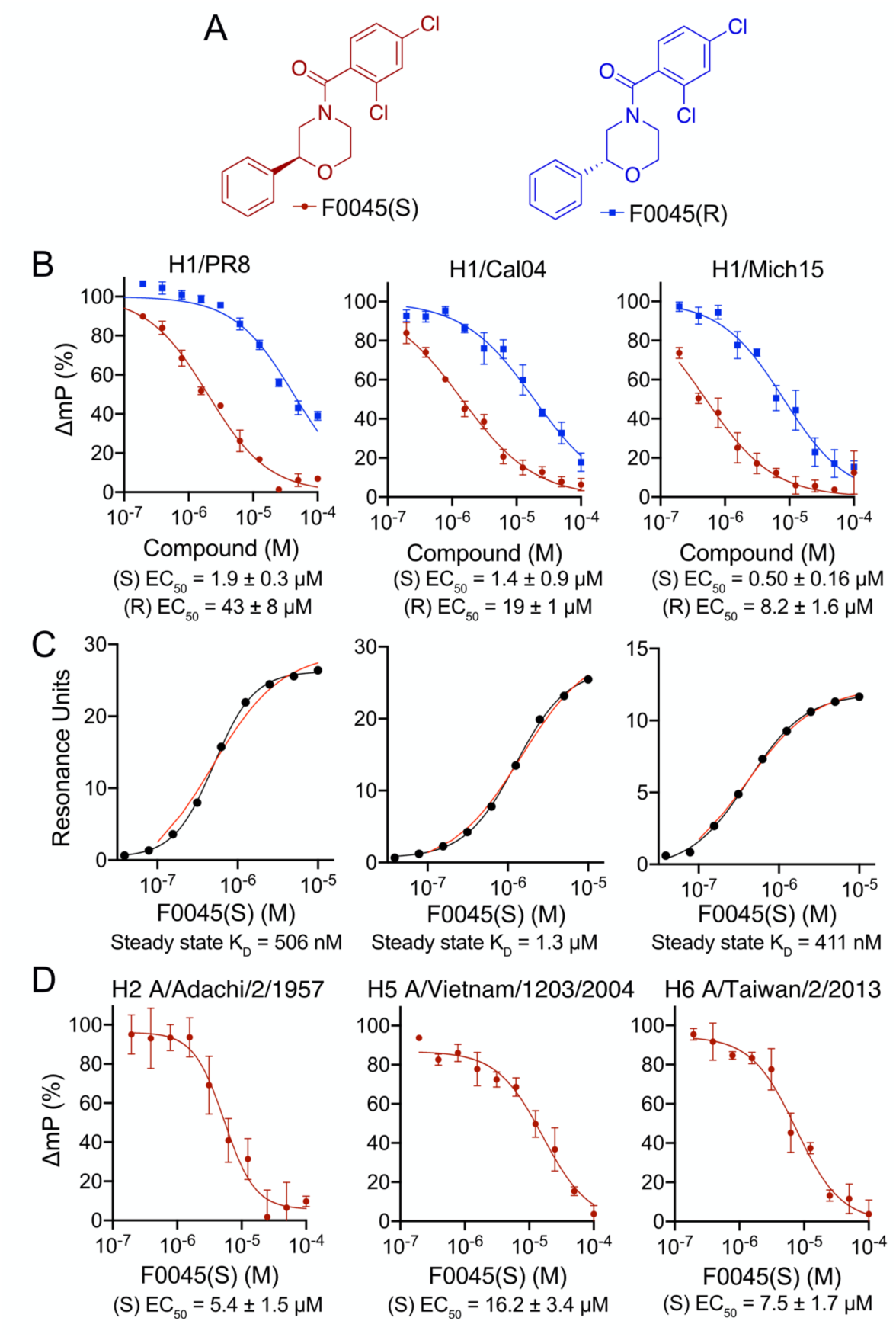
Characterization of F0045 enantiomers. (A) Chemical structures of F0045(S) and F0045(R). (B) Dose-response comparison (200 nM to 100 μM) of F0045(S) and F0045(R) binding a panel of H1 HAs as measured by FP P7-TAMRA probe competition assay. The assay consisted of a solution of PBS, pH 7.4, 0.01% Triton X-100) and 75 nM P7-TAMRA probe and H1 HA (30 nM H1/PR8; 30 nM H1/Cal04; 100 nM H1/Mich15) mixed for several seconds at room temperature prior to measuring FP. DMSO and 300 nM P7 peptide represent the negative and positive controls, respectively. FP was measured in triplicate. (C) Steady state kinetic dose responses of F0045(S) (200 nM to 250 μM) against H1 HAs (H1/PR8, H1/Cal04 and H1/Mich15) as measured by SPR. (D) Dose-response comparison of F0045(S) binding a panel of group 1 HAs (50 nM for H2 A/Adachi/2/1957 and H5 A/Vietnam/1203/2004; 55 nM for H6 A/Taiwan/2/2013) measured as in (A).

We then subjected these F0045 enantiomers and H1 HAs to additional biophysical methods to further validate and compare stem binding and affinity. For example, differential scanning fluorimetry (DSF) analysis confirmed the preference of H1/PR8, H1/Cal04, and H1/Mich15 HAs for F0045(S) over F0045(R), as F0045(S) stabilizes the HA protein from temperature-induced unfolding at much lower concentrations than F0045(R) (**Figs. S3, S4**). Additionally, incubation of either F0045 enantiomer with group 2 H3/HK68 HA did not result in any observable increase in melting temperature (**Fig. S4**). We anticipated group 2 HAs would have limited to no affinity for F0045 because the binding epitope has a few important differences that include an additional glycosylation site at Asn38 and some other amino-acid substitutions that, when combined, would hinder F0045 binding. Steady state and kinetic binding of F0045(S) and (R) to group 1 HAs were also measured by surface plasmon resonance (SPR). Steady state K_D_ values of F0045(S) measured by SPR correlate with the EC_50_ values obtained from the FP competition assay (**Fig. 3C**). With respect to kinetic binding, F0045(S) demonstrated lower K_D_ values against different H1 HAs in comparison to F0045(R): 0.3 μM (S) vs. 4.7 μM (R) for H1/PR8; 0.8 μM (S) vs. 14 μM (R) for H1/Cal04; and 0.5 μM (S) vs. 5.6 μM (R) for H1/Mich15 (**Fig. S5**). The improved K_D_ values for F0045(S) are primarily a result of a significantly reduced dissociation rate. For example, the k_off_ of F0045(S) vs. (R) for H1/PR8 was 0.03 s^-1^ and 0.18 s^-1^, respectively (**Fig. S5**).

We next wanted to determine the breadth of F0045(S) binding to other group 1 HAs. Using our P7-TAMRA FP assay, we measured relative EC_50_ values for H2 A/Adachi/2/1957 (EC_50_ = 5.4 ± 1.5 μM), H5 A/Vietnam/1203/2004 (EC_50_ = 16.2 ± 3.4 μM), and H6 A/Taiwan/2/2013 (EC_50_ = 7.5 ± 1.7 μM) (**Fig. 3D**). Interestingly, the observed trend of decreased affinities for other group 1 HAs is similar to the diminished affinities previously reported for stem-targeting molecule JNJ4796 (22). Future HTS efforts will focus on these individual HA subtypes to try to identify common small-molecule motifs that can be optimized in parallel across group 1 HAs.

We investigated the structural basis of binding and neutralization of influenza A virus by F0045(S). A co-crystal structure of F0045(S) in complex with H1/PR8 HA was determined at 2.69 Å resolution (**Fig. 4A, Table S1**). F0045(S) exhibited well-defined electron density and recognizes the hydrophobic cavity at the interface of the HA1-HA2 in the HA stem region (**Figs. 4A, S6**). This region consists of residues His^18^, His^38^-Leu^42^ and Thr^318^ from HA1, and Asp^19^, Trp^21^ and Glu^38^-IIe^56^ from helix-A of HA2. This HA1-HA2 interface consists of a number of small hydrophobic pockets, a few of which are occupied by the A- to C-rings of F0045(S) (**Fig. 4A**). Analysis of molecular interactions of F0045(S) with H1/PR8 HA demonstrated a series inter-molecular polar and nonpolar interactions (**Fig. 4B, S7**). The amide carbonyl of the A-ring of F0045(S) makes a direct hydrogen bond interaction with the sidechain hydroxyl of Thr^318^ from HA1, whereas Cγ2 CH from Thr^318^ makes a CH-π interaction with the A-ring. Similarly, the C-ring of F0045(S) makes CH-π interactions with His^18^ and Trp^21^ from HA1 and HA2, respectively (**Fig. 4B**). These C-ring interactions of F0045(S) are strikingly similar to the interactions made by the D-ring of small molecule JNJ4796 in the JNJ4796-H1 HA complex (PDB ID 6CF7) (**Fig. 4C**) (22). Nonpolar interactions of F0045(S) include contacts with the sidechains of Thr^318^, Val^40^ from HA1 and Trp^21^, Thr^41^, IIe^45^, IIe^48^ and Val^52^ from HA2. This network of polar and nonpolar interactions of F0045(S) with residues in the stem binding site appears then to stabilize the HA1/HA2 interface in its pre-fusion conformation and prevent pH-induced conformational changes to the post-fusion form (**Fig. 2C**).

**Figure 4.**
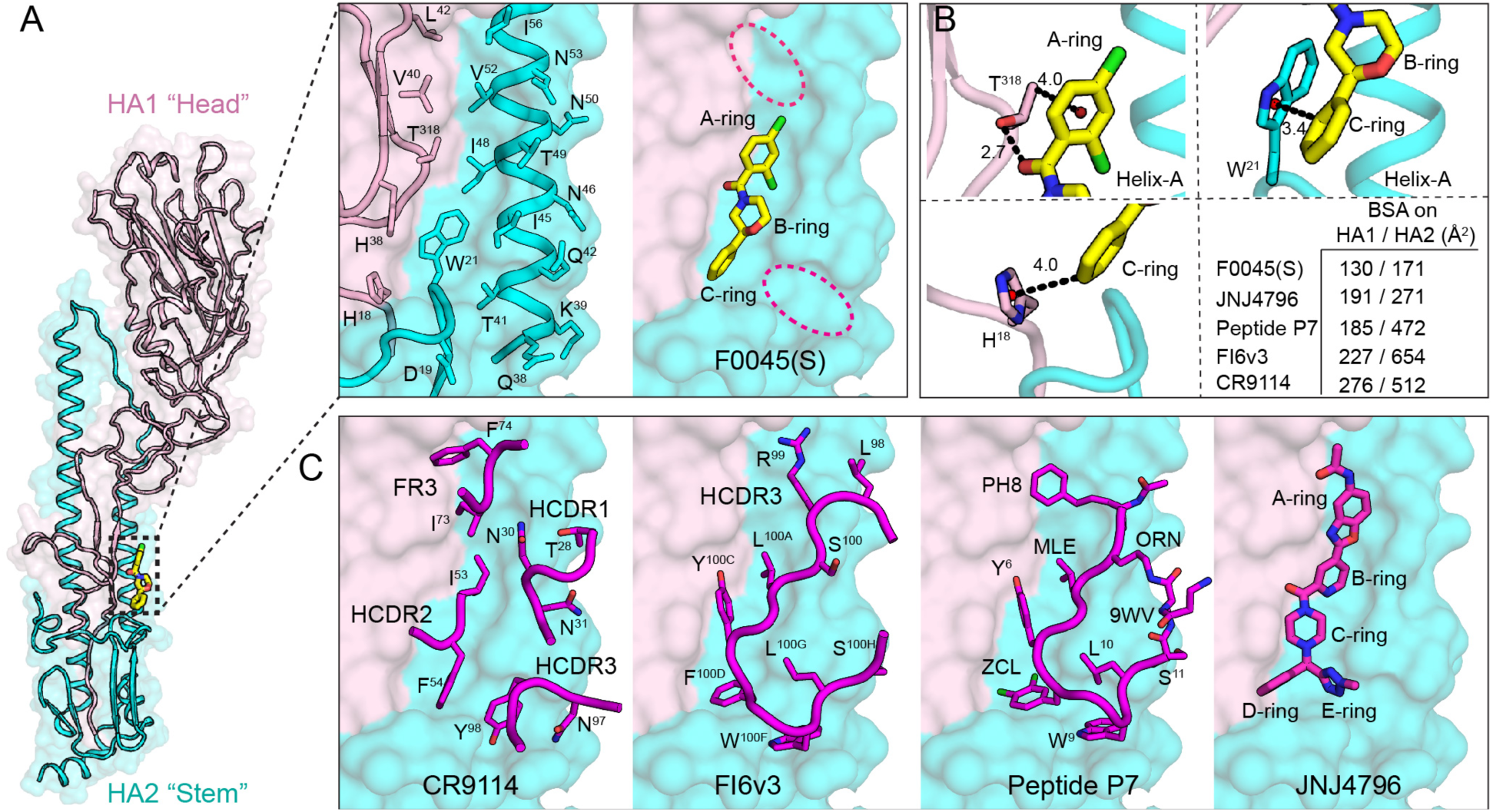
Structure and molecular interactions of small molecule F0045(S) with influenza A HA. (A) Crystal structure of influenza A HA from group 1 H1N1 A/Puerto Rico/8/1934 (H1/PR8) (HA1-light pink and HA2-cyan cartoon with molecular surface representation) in complex with small molecule F0045(S) (carbon, oxygen, nitrogen, and chlorine are represented in yellow, red, blue, and green, respectively). One protomer of the H1/PR8 trimer is represented. F0045(S) binding site residues on the molecular surface of H1/PR8 HA shown in sticks on a backbone tube representation. Additional hydrophobic pockets around F0045(S) are highlighted in dotted red ovals. (B) Molecular interaction in the F0045(S) - H1/PR8 HA complex. The centroids of the rings are shown in a red sphere, hydrogen bonds in black dotted lines, and distances are measured in angstroms (Å). Buried surface area (BSA) of F0045(S) and other inhibitors on HA. (C) Overlay of structures of F0045(S)-H1/PR8 with HA-interacting loop residues from the Fabs of HA-Fab complexes of CR9114 (PDB ID 4FQI), FI6v3 (PDB ID 3ZTN), and with cyclic peptide P7 (PDB ID 5W6T) and small molecule JNJ4796 (PDB ID 6CF7). Antibody CDR loops and cyclic peptide are represented as a backbone tube with side chains as sticks and JNJ4796 in stick representation, respectively.

Overall, F0045(S) buries ∼130 and 171 Å^2^ on HA1 and HA2, respectively, of H1/PR8 HA, which is a fairly small surface area compared to the known small molecule, peptide and antibodies targeting this region (**Fig. 4B,C**). F0045(S) occupies the same region as targeted by HCDR2 and HCDR3 loops of bnAbs CR9114 and FI6v3, respectively, and the antibody-inspired peptide P7. The binding mode of F0045(S) is also strikingly similar to the binding mode of B-, C- and D-rings of small molecule JNJ4796 (**Fig. 4C**). Thus, with a minimal footprint as compared to the currently known inhibitors and the presence of surrounding additional unoccupied pockets in the binding site, F0045(S) presents an excellent opportunity to elaborate the small molecule into nearby regions to improve the binding and neutralization activity against influenza viruses (**Fig. 4A-C**).

To elucidate structural basis for the stereospecific binding of F0045 towards group 1 influenza A HAs. We compared the two different configurations of F0045 in the binding site of H1/PR8. Based on the crystal structure of H1/PR8 with F0045(S) (**Fig. 5A**), we inverted the S-stereocenter in the B-ring of F0045(S) to R-form and compared the binding mode of these two different configurations that affect the relative disposition of the C-ring. Compared to the C-ring of F0045(S) in F0045(S)-H1/PR8 complex, the modeled F0045(R) shows that the C-ring would now be located inward with increased proximity to helix-A and further steric clash with HA2 IIe^45^ and Trp^21^ (**Fig. 5B**). These steric considerations could explain the reduced binding and neutralization activity of F0045(R) against group 1 HAs as compared to F0045(S) (**Fig. 5A**).

**Figure 5.**
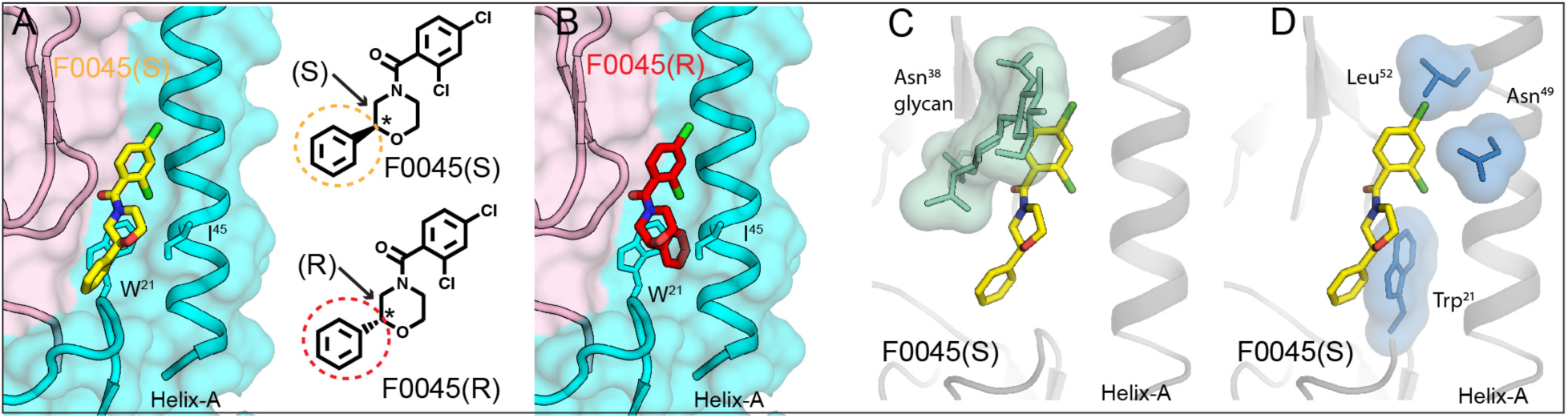
Structural basis of stereoselectivity and group 1 HA specificity. (A) S-configuration of F0045 (yellow stick) in complex H1/PR8 HA. (B) Modeled R-configuration of F0045 (red stick) in complex with H1/PR8 HA. HA1 is in light pink and HA2 in cyan. HA2 residues Trp^21^ and Ile^45^ are shown in stick representation. S- and R-configurations of the C-ring of F0045 are highlighted in yellow and red circles, respectively. Asterisk indicates the stereocenter. (C and D) Superimposition of F0045(S) (yellow stick) bound to H1/PR8 HA on apo H3/HK68 (PDB ID 4FNK) (HA is a gray cartoon with selected residues in green and blue sticks with their surface representation). Only F0045(S) from the H1/PR8 HA complex is shown.

Further, to understand group 1 HA binding specificity of F0045(S), the structure of F0045(S) bound to H1/PR8 HA was superimposed with the group 2 apo H3/HK68 HA structure (PDB ID 4FNK) (**Fig. 5C,D**). The key differences in the F0045(S) binding site on H1/PR8 vs apo H3/HK68 HAs are presence of a glycosylation site at HA1 Asn^38^ in group 2 HA, the orientation of HA2 Trp^21^, and the presence of Asn^49^ and Leu^52^ in helix-A of group 2 H3/HK68 HA that could all lead to steric hindrance of F0045(S) with group 2 HAs and render it group 1 HA specificity. This observation is consistent with group 1 specificity of antibody CR6261 (30), cyclic peptide P7 (21), and small molecule JNJ4796 (22).

The most stringent *in vitro* analysis to assess the potential antiviral activity of chemical and/or biological interventions is with cell-based infection assays. MDCK-SIAT1 cells die after 72 hrs of incubation with influenza virus, as previously described (22, 31, 32). Increasing concentrations (976 nM to 500 μM) of F0045(S) and F0045(R) were co-incubated with MDCK-SIAT1 cells, respectively. Small molecules were added simultaneously with influenza virus (50 TCID_50_, median tissue culture infectious dose), or alone to assess toxicity, and incubated with MDCK-SIAT1 cells for 72 hr at 37 °C in 5% CO_2_ incubator. Cellular viability was then quantified using CellTiter-Glo^®^ and luminescence measured on a PerkinElmer EnVision plate reader. We showed F0045(S) and (R) are toxic to MDCK-SIAT1 cells with cytotoxic concentration (CC)_50_ values >130 μM (**Fig. 6A**); however, co-incubation with an influenza virus expressing the HA from H1N1 A/Puerto Rico/8/1934 (22) demonstrated F0045(S) selectively protects MDCK-SIAT1 cells from virus-induced cell death with EC_50_ = 100 ± 4 μM. Conversely, no observable virus neutralization was evident from the presence of F0045(R) consistent with the selectivity of H1 virus for the S-enantiomer (**Fig. 6B**). We next tested the ability of F0045(S) to prevent infection with H1N1 viruses expressing A/Beijing/262/1995 (H1/Beijing), H1/CA04, and A/Solomon Islands/3/2006 (H1/SI06), as well as H5N1 pseudovirus expressing H5 A/Vietnam/1203/2004. F0045(S) readily neutralized MDCK-SIAT1 infection by H1/Beijing (EC_50_ = 1.6 ± 0.1 μM), H1/Cal04 (EC_50_ = 3.9 ± 2.1 μM), and H5 A/Vietnam/1203/2004 (EC_50_ = 22.8 ± 2.3 μM) (**Fig. 6C**); however, F0045(S) did not protect cells from infection by H1/SI06-expressing virus. One possible explanation for the lack of protection is attributable to the fusion stability of each HA. For example, viruses can escape the neutralization effects of HA stem-binding bnAbs by altering their HA composition such that more acidic conditions are necessary for fusion (33-35). Theoretical pI values of the HAs vs. measured EC_50_ values showed that H1/SI06 has the lowest pI as well as the most unfavorable compound neutralization capability (**Fig. S8**).

**Figure 6.**
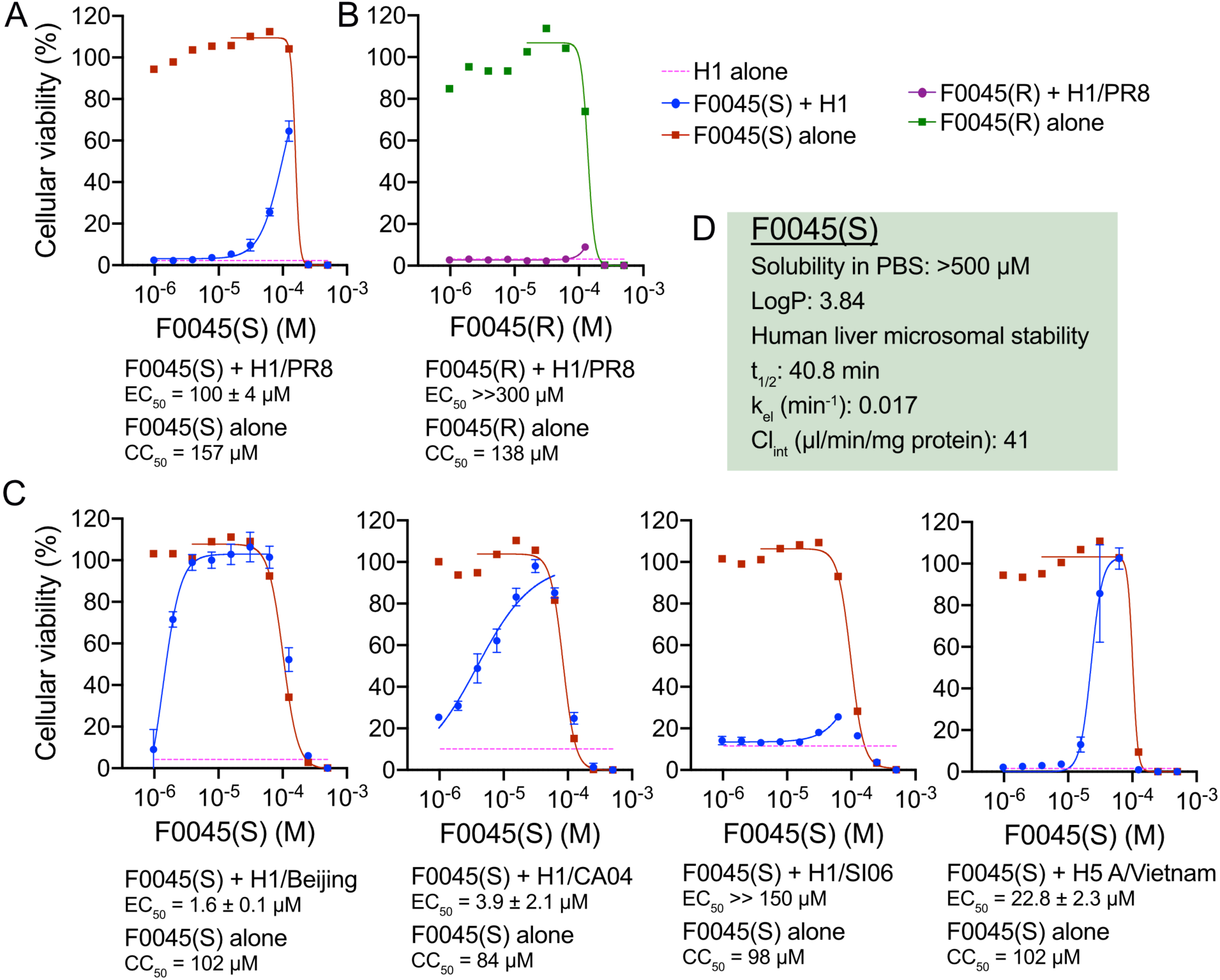
Cell-death assay of MDCK-SIAT1 cells. (A) Incubation of H1/PR8 influenza virus with F0045(S) measured at 72 h. F0045(S) protects cells from cell death at higher concentrations (EC_50_ = 100 ± 4 μM) and at concentrations lower than the measured CC_50_ of the compound (red squares). (B) F0045(R) does not protect MDCK-SIAT1 from H1/PR8-induced cell death (purple circles). F0045(R) has a similar toxicity to the cells as the S-enantiomer (green squares). (C) Virus neutralization assay with dose-response of F0045(S) against H1/Beijing, H1/Cal04, H1/SI06, and H5 A/Vietnam/1203/2004. (D) Drug-likeness of F0045(S): LogP is predicted with ChemDraw Ultra 19.0, human liver microsomal stability performed as described (45) with the elimination constant (k_el_), half-life (t_1/2_), and intrinsic clearance (Cl_int_) determined by the plot of ln(area-under-the-curve) versus time, using linear regression analysis.

In conclusion, we designed a robust and thorough FP binding assay compatible with high-throughput screening to identify molecules with affinity to the highly conserved stem region of influenza A group 1 HAs. The success of identifying a new HA inhibitor from a commercially available small-molecule library that neutralizes influenza in vitro and protects host cells from apoptosis demonstrates the advantages of our simple, low-cost, and efficient assay. Our small molecule hit differs from the previous JNJ4796 stem-targeting molecule that has been extensively optimized for *in vitro* and *in vivo* pharmacokinetics. For example, F0045(S) is less toxic to human cells (*e.g.*, CC_50_ > 130 μM for F0045(S) vs. >10 μM for JNJ4796) and represents an excellent lead molecule for medicinal chemistry, as the compound has reasonable stability in human microsome assays (t_1/2_ = 40.8 min (**Fig. 6D**)). We are actively improving the affinity and breadth of F0045(S) with iterative rounds of structure-based design and synthesis of structure-activity relationship libraries. While group 1 HAs have a known high-affinity peptide binder that can be modified with a fluorophore, additional methodologies and probes are urgently needed to interrogate group 2 HAs for new small-molecule inhibitors. We posit that similar strategies of mimicking bnAb CDR stem-targeted loops, such as the scaffold used for the P7 peptide design, will open up new avenues to explore all types, subtypes and lineages of HAs. Our ultimate goal is to discover cost-effective novel HA-based small molecules that neutralize all pandemic, seasonal, and emerging influenza A and B viruses.

## Materials and Methods

### P7-TAMRA probe synthesis

P7 was prepared in a similar way as previously reported (21). Briefly, the linear peptide was synthesized on a 0.3-mmol scale using 2-chlorotrityl chloride resin by manual Fmoc-based SPPS. Lactam cyclization of the crude linear peptide was performed at high dilution in DMF, with 3 eq. of HBTU, HOBt and DIPEA and the peptide was purified by reverse-phase HPLC. The product was characterized by LC-MS and subsequently reacted in 1 mL DMSO with 1.2 eq TAMRA succinimidyl ester (5(6)-TAMRA, SE) in the presence of 3 eq. DIPEA. The product was purified by reverse-phase HPLC and characterized by LC-MS. C_104_H_126_Cl_2_N_17_O_23_ [M^+^H]^+^: 2053.14, found (1)^+^=1026.92 (**Figs. 1B, S9**). Please see Supporting Information for full synthetic procedures.

### Expression and purification of the influenza A hemagglutinin

The HAs used for binding and crystallization studies were expressed using baculovirus expression system as described previously (36). Please see Supporting Information for details regarding procedures.

### Polarization assay

P7-TAMRA probe was incubated at a final concentration of 75 nM in the presence of group 1 HA trimer (30 nM final concentration for H1/PR8 and H1/Cal04; 100 nM for H1/Mich15; 50 nM for H2 A/Adachi/2/1957 and H5 A/Vietnam/1203/2004; 55 nM for H6 A/Taiwan/2/2013) in an assay buffer containing PBS, pH 7.4 and 0.01% Triton X-100. A 100 μL volume of P7-TAMRA probe and HA was dispensed into a black 96-well Costar flat-bottom polystyrene plate prior to FP measurement. Dose-dependent competition assays to determine relative EC_50_ values of P7, bnAb S9-3-37, F0045(S) and (R), DMSO or aqueous stock solutions were added to the pre-mixed P7-TAMRA probe and HA, vortexed for 10 sec at 1000 rpm with FP immediately read on a PerkinElmer EnVision plate reader. All assay conditions required n > 3 replicates. Data were analyzed using GraphPad Prism to determine EC_50_.

### High-throughput screen

10 µL of solution containing 30 nM H1/PR8 HA and 75 nM P7-TAMRA probe in assay buffer (PBS, pH 7.4 and 0.01% Triton X-100) was added into each well of a black 384-well Greiner low volume plate with a Thermo Multidrop 384 dispenser. Next, 100 nL library compounds (2 mM stock) were added into each well using a Biomek FXP Laboratory Automation Workstation, and each plate incubated at room temperature for 30 mins. Fluorescence polarization was then measured on a PerkinElmer EnVision plate reader (ex. filter: 531 nm; em. filter: 595p and 595s; mirror: BODIPY TMR dual). Vehicle DMSO and 300 nM P7 peptide served as the negative and positive controls, respectively, and represented the upper and lower FP values for normalization of ΔmP.

### Trypsin susceptibility (TS) assay

5 μM H1/PR8 HA was pre-incubated with 50 μM of P7 peptide, P7-TAMRA probe, or F0045 for 30 min at room temperature (control reactions consisted of 2% DMSO vehicle). The pH of each reaction was lowered using 1 M sodium acetate buffer (pH 5.0). One reaction was retained at pH 7.4 to assess digestion at neutral pH. The reaction solutions were then thoroughly mixed and incubated for 20 min at 37 °C. The solutions were subsequently equilibrated to room temperature and the pH was neutralized by addition of 200 mM Tris buffer, pH 8.5. Trypsin-ultra™ (NEB Inc.) was added to all samples at final ratio of 1:50 by mass and the samples were digested for 30 min at 37 °C. After incubation with trypsin, the reactions were equilibrated to room temperature and quenched by addition of non-reducing SDS buffer and boiled for ∼2 min at 100 °C. All samples were analyzed by 4-20% SDS-PAGE gel and imaged using BioRad ChemDoc™ imaging system.

### Crystallization and structure determination of F0045(S)-H1/PR8 HA complex

Gel filtration fractions containing H1/PR8 HA was concentrated to ∼10 mg/mL in 20mM Tris, pH 8.0 and 150mM NaCl. Before setting up crystallization trials, F0045(S) at ∼5 molar excess was incubated with H1/PR8 HA for ∼30 minutes at room temperature and centrifuged at 10,000g for ∼4-5 minutes. Crystallization screens were set up using the sitting drop vapor diffusion method using our automated CrystalMation robotic system (Rigaku) at TSRI. Within 3-7 days, diffraction-quality crystals were obtained using 0.2 M magnesium nitrate and 20% w/v PEG3350 as precipitant at 4 °C. Crystals were cryoprotected with 5-15% ethylene glycol, and then flash cooled and stored in liquid nitrogen until data collection. Diffraction data were processed with HKL-2000 (37). Initial phases were determined by molecular replacement using Phaser (38) with an HA model from H1/PR8 (PDB ID 5W5S). Refinement was carried out in Phenix (39), alternating with manual rebuilding and adjustment in COOT (40). Detailed data collection and refinement statistics are summarized in **Table S1**.

### Structural analyses

Two-dimensional depiction of the F0045(S) binding sites on HA rendered using the Flatland Ligand Environment View (FLEV) mode of Lidia module in COOT (40). Surface areas buried on the H1 PR8 HA upon binding of F0045(S) was calculated with the Protein Interfaces, Surfaces and Assemblies (PISA) server at the European Bioinformatics Institute (41). Fab FI6v3 (PDB ID 3ZTN), designed peptide P7 (PDB ID 5W6T) and small molecule JNJ4796 (PDB ID 6CF7) bound to an H1N1 HA, and Fab CR9114 (PDB ID 4FQI) bound to H5N1 HA were used for buried surface area calculations. MacPyMol (DeLano Scientific) was used to render structure figures. The final coordinates were validated using MolProbity (42).

### Accession code

The coordinates and structure factors for the F0045(S) with H1/PR8 HA have been deposited with the Protein Data Bank under accession code 6WCR.

### Virus neutralization assay (VNA)

25,000 MDCK-SIAT cells (Madin-Darby canine kidney cells overexpressing the α-2,6-linked sialic acid receptor) were plated into each well of a 96-well plate in a total volume of 100 μl DMEM supplemented with 1x penicillin-streptomycin, 10% FBS (fetal bovine serum), and 1x NEAA (non-essential amino acids). Plates were permitted to incubate overnight at 37 °C, 5% CO_2_ incubator for MDCK-SIAT1 cells to attach to the plate. Cells were washed twice with D-PBS and medium replaced with 100 μl OptiMEM diluent containing 0.8 μg/ml N-tosyl-L-phenylalanine chloromethyl ketone (TPCK)-treated trypsin and 0.5% DMSO. Four H1N1 influenza A viruses (H1/PR8, H1/Beijing, H1/SI06, and H1/Cal04) and one H5N1 influenza A pseudovirus (H5 A/Vietnam/1203/2004) were used in this assay (43, 44). 500 μM of F0045(S)/(R) in 0.5% DMSO were 2-fold serially diluted in 50 TCID_50_ virus diluent in triplicates and incubated with cells at 37 °C for 72 hr in a 5% CO_2_ incubator. CellTiter-Glo^®^ luminescent cell viability reagent was then added in each well according the manufacturer’s instructions. Luminescence was measured on a PerkinElmer EnVision plate reader and EC_50_ values were calculated with GraphPad Prism (n = 3 for each condition).

## Author Contributions

I.A.W. and D.W.W. designed the research; Y.Y. performed synthesis, HTS assay, and compound validation; R.U.K. purified protein and performed the x-ray structure studies; Y.Y. and J.L.W. performed surface plasmon resonance; Y.Y. and S.K. performed differential scanning fluorimetry; R.U.K. and X.Z. performed trypsin-susceptibility assay; C-C.D.L., N.C.W., and D.W.W. performed cellular infection assays; Y.Y., R.U.K., C-C.D.L, N.C.W., I.A.W., and D.W.W. analyzed the data; Y.Y., R.U.K., I.A.W., and D.W.W. composed the paper and all authors edited and approved its contents.

The authors declare no conflict of interest.

## Acknowledgments

We thank H. Rosen, R.L. Wiseman, and L.L. Lairson for access to instrumentation. This work was supported by The Scripps Research Institute (D.W.W.), National Institutes of Health (NIH) R56 AI127371 (I.A.W. and D.W.W.), NIH K99 AI139445 (N.C.W.), and the Bill and Melinda Gates Foundation OPP1170236 (I.A.W.). C-C.D.L. was sponsored by an Academia Sinica Fellowship, Academia Sinica, Taiwan.

