## Supporting Information for "An influenza A hemagglutinin small-molecule fusion inhibitor identified by a new high-throughput fluorescence polarization screen"

### Supplementary Information Text

#### Materials and reagents

Fmoc-protected amino acids were purchased from Combi-Blocks and Oakwood Products, 5(6)-TAMRA, SE [5-(and-6)-carboxytetramethylrhodamine, succinimidyl ester] from AAT Bioquest, and (S)-2-phenylmorpholine and (R)-2-phenylmorpholine from Astatech. Greiner Bio-One 384 wells and pyrrolidine were purchased from Fisher Scientific. CellTiter-Glo® luminescent cell viability assay reagent was purchased from Promega Corp. All other reagents were of analytical grade and used without further purification. Milli-Q water was used in all experiments unless otherwise stated.

#### Synthesis

**P7 peptide.** P7 was prepared in a similar way as previously reported (1). The linear peptide was synthesized on a 0.3 mmol scale using 2-chlorotrityl chloride resin by manual Fmoc solid phase peptide synthesis. The first amino acid was loaded with an anhydrous DCM solution of 3 eq. Dde-L-Dap(Fmoc)-OH in the presence of 5 eq. N,N-diisopropylethylamine (DIPEA). The resin was then capped with freshly made solution of DCM/MeOH/DIPEA (17:2:1) for 1 hour. The N-Dde protecting group was then removed by reaction with a 2% hydrazine hydrate solution in DMF, followed by N-Boc-β-alanine coupling. Standard Fmoc deprotection (20% Pyrrolidine in DMF) and coupling protocols (3 eq. Fmoc amino acid, HCTU and DIPEA) were used until the linear peptide was synthesized. The protected linear peptide was then cleaved from the resin using 20% hexafluoroisopropanol in DCM.

Finally, the lactam cyclization of crude linear peptide was performed at high dilution in DMF, with 3 eq. of HBTU, HObt and DIPEA. The reaction was performed at room temperature for 2 days, followed by full side-chain deprotection with solution of TFA/triisopropylsilyl/$H_2O$ (95:2.5:2.5). The final product was purified by reverse-phase HPLC (linear gradient from 90% water and 10% MeCN to 100% MeCN). The product was characterized by LC-MS. Calc'd for $C_{79}H_{106}Cl_2N_{15}O_{19}$ $[M+H]^+$: 1640.70, found 1641.12 (**Fig. S9**).

**P7-TAMRA probe.** 1 eq. P7 peptide and 1.2 eq TAMRA succinimidyl ester (5(6)-TAMRA, SE) were dissolved in 1mL DMSO and 3 eq. DIPEA was then added to the solution. The solution was stirred in the dark at room temperature overnight. Water was then added to quench the reaction and purified by reverse-phase HPLC with 95% yield (linear gradient from 90% water and 10% MeCN to 100% MeCN). The product was characterized by LC-MS. Calc'd for $C_{104}H_{126}Cl_2N_{17}O_{23}$ $[M+H]^+$: 2053.14, found $(1)^+$=1026.92 (**Fig. S9**).

**F0045(S).** EDCI·HCl (264 mg, 1.38 mmol) and DIPEA (410 µl, 2.76 mmol) were added to a solution of (S)-2-phenylmorpholine (150 mg, 0.92 mmol), 2,4-dichlorobenzoic acid (193 mg, 1.01mmol) and HOBt (186 mg, 1.38 mmol) in DCM, and stirred for 24 hr at room temperature under nitrogen. The solvent was removed *in vacuo* and the residue purified by column chromatography on silica gel eluting with DCM:MeOH=100:1 to 100:5 (by volume), to furnish the title compound as a white foam (95% yield). $^1$H NMR (500 MHz, $CDCl_3$) δ 7.52-7.18 (m, 8H, ArH), 4.89-4.66 (m, 1H, CH), 4.65-2.88 (m, 6H, $CH_2$). 13C NMR (500 MHz $CDCl_3$) δ 166.26, 138.75, 135.89, 134.05, 131.36, 129.77, 129.15, 128.64, 128.33, 127.92, 126.18, 78.54, 66.91, 53.65, 47.91. MS (ESI). Calc'd for $C_{17}H_{16}Cl_2NO_2$ $[M+H]^+$: 336.05, found 336.13 (**Figs. S10, S11**).

**F0045(R).** EDCI·HCl (264 mg, 1.38 mmol) and DIPEA (410 µl, 2.76 mmol) were added to a solution of (R)-2-phenylmorpholine (150 mg, 0.92 mmol), 2,4-dichlorobenzoic acid (193 mg, 1.01 mmol) and HOBt (186 mg, 1.38 mmol) in DCM, and stirred for 24 hours at room temperature under nitrogen. The solvent was removed *in vacuo* and the residue purified by column chromatography on silica gel eluting with DCM:MeOH=100:1 to 100:5 (by volume), to furnish the title compound as a white foam (95% yield). $_1$H NMR (500 MHz, $CDCl_3$) δ 7.52-7.18 (m, 8H, ArH), 4.89-4.66 (m, 1H, CH), 4.65-2.88 (m, 6H, $CH_2$). 13C NMR (500 MHz $CDCl_3$) δ 166.26, 138.75, 135.89, 134.05, 131.36, 129.77, 129.15, 128.64, 128.33, 127.92, 126.18, 78.54, 66.91, 53.65, 47.91. MS (ESI). Calc'd for $C_{17}H_{16}Cl_2NO_2$ $[M+H]^+$: 336.05, found 336.16 (**Figs. S10, S11**).

**Methods**

**LC-MS characterization.** Small molecules or peptides were dissolved in a $H_2O$/MeCN solution mixture, and an aliquot was taken for LC-MS analysis. Separation was achieved by gradient elution from 10-100% MeCN in water (constant 0.1 vol% formic acid) over 12 min, isocratic with 100% MeCN from 12 to 17 min and returned to initial conditions and equilibrated for 3 min. The LC chromatograms were recorded by monitoring absorption at 220 and 280 nm.

**Expression and purification of influenza A hemagglutinin.** The hemagglutinin (HA) used for binding and crystallization studies were expressed using our baculovirus expression system as described previously (2). Briefly, each HA was fused with a gp67 signal peptide at the N-terminus and to a BirA biotinylation site, thrombin cleavage site, foldon trimerization domain and $His_6$-tag at the C-terminus. Expressed HAs were purified using metal affinity chromatography using Ni-NTA resin. For binding studies, each HA was biotinylated using BirA and purified by gel filtration chromatography. For crystallization trials, the HA was digested with trypsin (New England Biolabs) to produce uniformly cleaved (HA1/HA2), and to remove the trimerization domain and $His_6$-tag. The digested material was purified by gel filtration using Superdex 200 16/90 column on an ÄKTA (GE Healthcare Life Sciences).

**Surface plasmon resonance (SPR).** Direct binding of each compound to HA was measured by SPR using a Biacore 8K instrument (GE Healthcare) at 25 °C. Sensor chip SA (GE Healthcare) were used to create H1 biosensors by standard biotin-streptavidin coupling. 50 μg/ml HA diluted in PBS, pH 7.4 was injected for 15 min at 5 μl/min over a single flow cell. In total, six separate biotin-streptavidin-coupled H1 surfaces were generated during the course of this study with the following immobilization levels: H1/PR8 (5715 and 5042), H1/Mich15 (3081 and 3032), and H1/Cal04 (7547 and 8300) resonance units (RUs). All experiments were performed in a running buffer of PBS, pH 7.4, 0.005% (v/v) Tween-20, and 5% (v/v) DMSO (PBST-DMSO) using a flow rate of 30 μl/min. Solvent correction curves were obtained at the beginning and end by injecting varying DMSO concentrations (4.4, 4.6, 4.8, 5.0, 5.2, 5.4, 5.6, and 5.8% [v/v]). A reference flow cell was created on each sensor chip by injection of biotin.

To evaluate H1 binding, we diluted each compound from a 100 mM stock solution in neat DMSO to 20X of the desired concentrations, followed by dilution in a buffer containing 1.05X PBST to match the running buffer. Samples were injected in a dose-dependent manner over the H1 biosensor for 30 sec followed by 180 sec of dissociation phase and a subsequent wash step with a 50% (v/v) DMSO solution. The resulting sensorgrams were analyzed using Biacore 8K Evaluation Software as follows. Solvent correction curves were generated and applied to all datasets, and each sensorgram was double referenced by subtracting the nearest buffer blank injection. The signal just before injection stop of these corrected sensorgrams was treated as the H1-binding response for steady-state affinity analysis and fit using four-parameter variable slope nonlinear curve fitting in GraphPad Prism8 software.

**NanoDSF.** 10 μL of 0.2 mg/ml of HA protein in PBS, pH 7.4 was mixed with different concentrations of the sample (F0045(S), F0045(R)). DMSO and 3 μM P7 peptide were used as negative and positive controls. The mixture was then transferred to capillary and scanned in nanoDSF from 20 °C to 95 °C with 1 °C/min rate for protein unfold temperature.

***In vitro* human microsomal stability measurements.** Microsomal incubations were carried out in 96-well plates in 5 aliquots of 40 µL each (one for each time point). Liver microsomal incubation medium comprised of PBS (100 mM, pH 7.4), $MgCl_2$ (3.3 mM), NADPH (3 mM), glucose-6-phosphate (5.3 mM), glucose-6-phosphate dehydrogenase (0.67 units/ml) with 0.42 mg of liver microsomal protein per ml. In the control reactions the NADPH-cofactor system was substituted with PBS. Test compounds (2 μM, final solvent concentration 1.6%) were incubated with microsomes at 37 °C, shaking at 100 rpm. Each reaction was performed in duplicates. Five time points over 40 minutes were analyzed. The reactions were stopped by adding 12 volumes of 90% acetonitrile-water to incubation aliquots, followed by protein sedimentation by centrifuging at 5500 rpm for 3 min. Each reaction was performed in duplicate. Supernatants were analyzed using the HPLC system coupled with tandem mass spectrometer API 3000 (PE Sciex). The TurboIonSpray ion source was used in positive ion mode. Acquisition and analysis of the data were performed using Analyst 1.5.2 software (PE Sciex). The elimination constant ($k_{el}$), half-life ($t_{1/2}$) and intrinsic clearance ($Cl_{int}$) were determined by the plot of ln(area-under-the-curve) versus time, using linear regression analysis.

**Statistical analysis.** All data are shown as mean ± SD with $n \geq 3$.

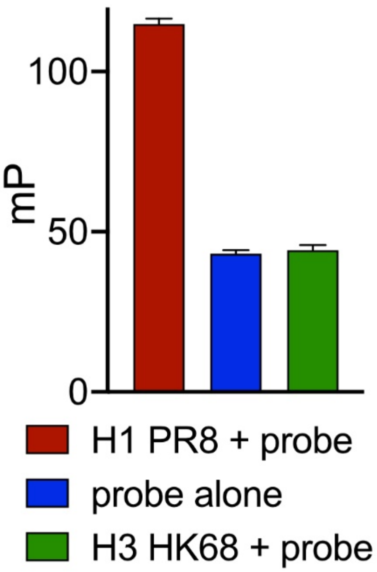


**Fig. S1.** FP assay and P7-TAMRA probe is specific for group 1 H1 versus group 2 H3 HA. 30 nM HA and 75 nM P7-TAMRA probe were mixed in PBS buffer (pH 7.4, 0.01% Triton X-100) for a few seconds. FP was then measured in 96-well plate by PerkinElmer EnVision plate reader at room temperature.

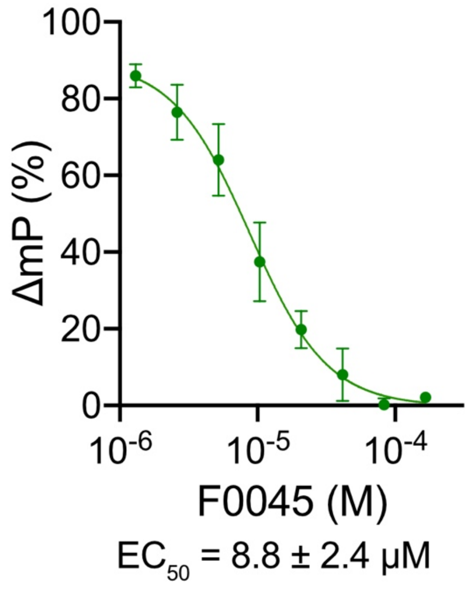


**Fig. S2.** P7-TAMRA probe (75 nM) and H1/PR8 HA (30 nM) competition assay against a dose-dependent increase of a racemic mixture of F0045.

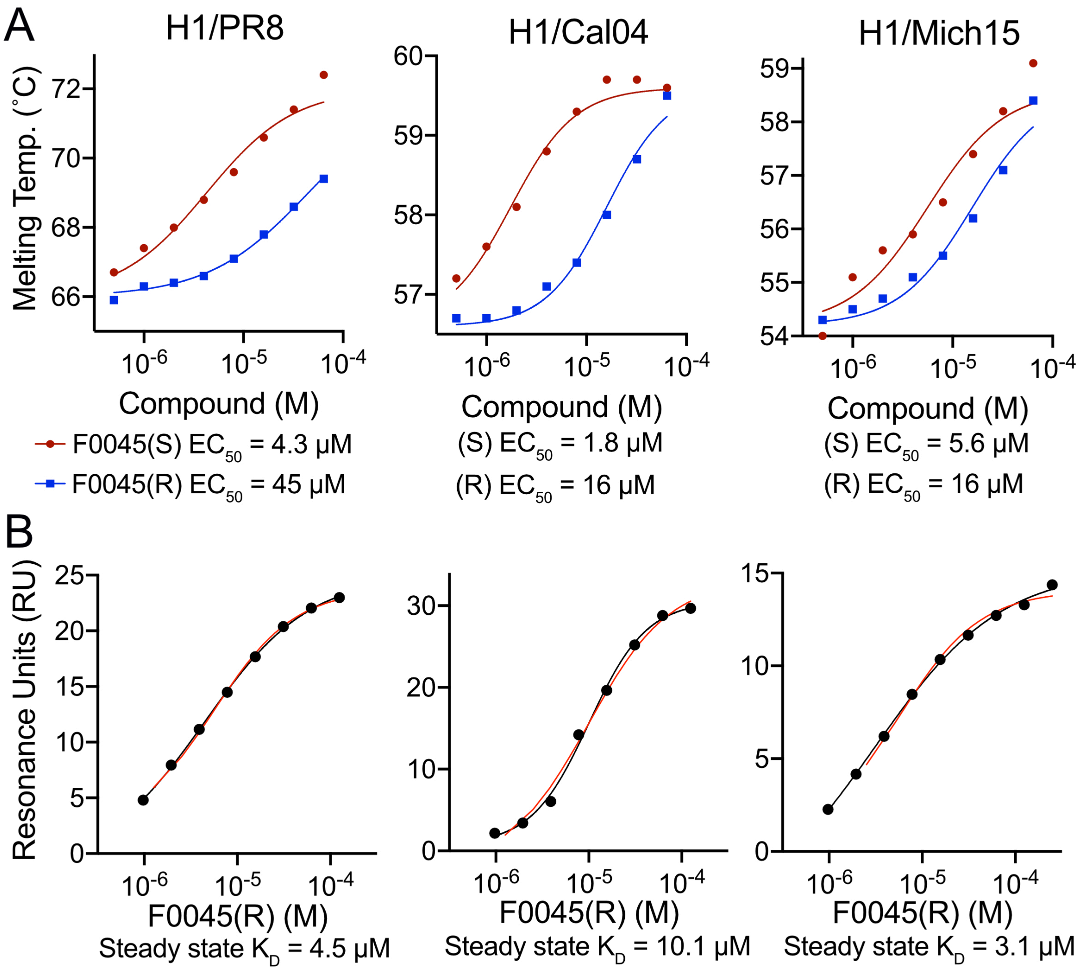


**Fig. S3.** Dose response of F0045(S) and (R) on temperature stability of H1 HAs and SPR of F0045(R) binding to HA**.** (A) Dose-response of F0045(S) vs. F0045(R) against different H1 HAs by DSF. F0045(S) and F0045(R) ranging from 0.5 μM to 64 μM were pre-mixed with 0.2 mg/ml HAs (H1/PR8, H1/Cal04, and H1/Mich15) in PBS buffer (pH 7.4), and HA protein unfolding temperature was then measured by nanoDSF. (B) Dose response of F0045(R) against different H1 HAs (H1/PR8, H1/Cal04, and H1/Mich15) by SPR. Different concentrations (250 μM to 200 nM) of F0045(R) against resonance units (RU) of different H1 HAs are plotted to determine the steady-state $K_D$.

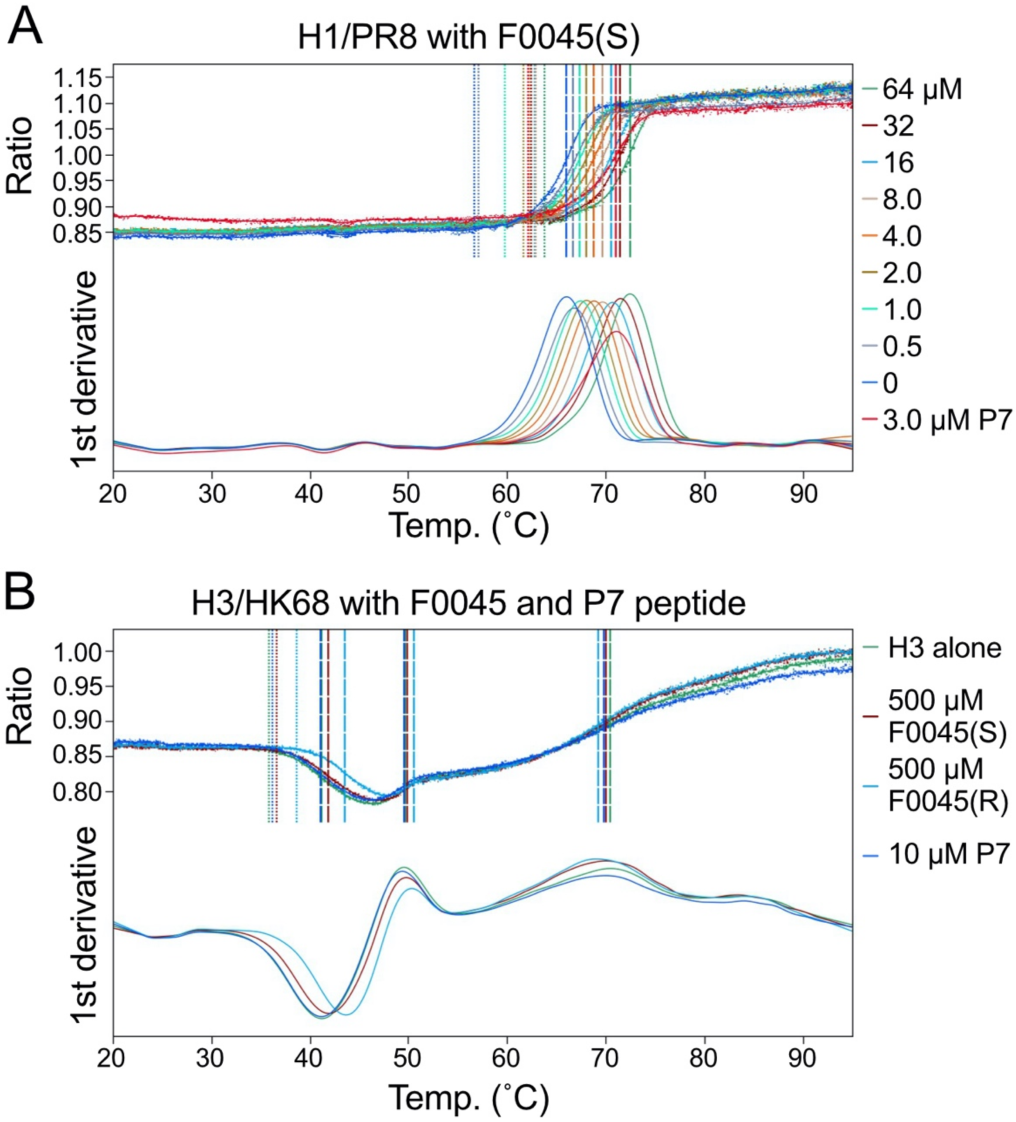


**Fig. S4**. DSF melting curves (A) H1/PR8 HA with increasing concentrations of F0045(S) (0 – 64 μM). (B) Melting temperature of H3/HK68 HA as control does not change in the presence of 10 μM P7, 500 μM F0045(S), or 500 μM F0045(R).

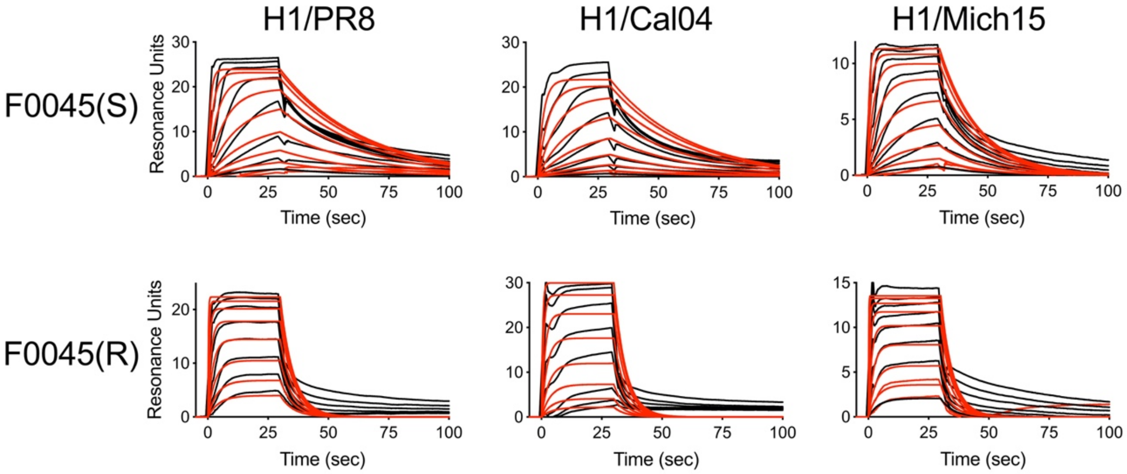


| | H1/PR8-Biotin | | | | |
|---|---|---|---|---|---|
| | Ka (1/Ms) | Koff (1/s) | Kinetic KD (M) | Rmax (RU) | Chi2 (RU2) |
| F0045(R) | 3.88E+04 | 1.84E-01 | 4.73E-06 | 23.2 | 1.66 |
| F0045(S) | 8.99E+04 | 2.91E-02 | 3.24E-07 | 24.7 | 1.8 |

| | H1/Mich15-Biotin | | | | |
|---|---|---|---|---|---|
| | Ka (1/Ms) | Koff (1/s) | Kinetic KD (M) | Rmax (RU) | Chi2 (RU2) |
| F0045(R) | 3.62E+04 | 2.02E-01 | 5.57E-06 | 13.8 | 7.28E-01 |
| F0045(S) | 1.29E+05 | 6.08E-02 | 4.72E-07 | 11.8 | 1.96E-01 |

| | H1/CA04-Biotin | | | | |
|---|---|---|---|---|---|
| | Ka (1/Ms) | Koff (1/s) | Kinetic KD (M) | Rmax (RU) | Chi2 (RU2) |
| F0045(R) | 1.48E+04 | 2.07E-01 | 1.39E-05 | 33.3 | 4.09 |
| F0045(S) | 4.05E+04 | 3.10E-02 | 7.65E-07 | 23.3 | 2.88 |

**Fig. S5**. SPR sensorgrams for HA-F0045(R)/(S) binding. Different concentrations of F0045(S) and F0045(R) were passed over immobilized HAs (H1/PR8, H1/Cal04 and H1/Mich15). Representative sensorgrams in resonance units (RU) plotted against time of injection are shown. Black lines are the experimental trace obtained from SPR experiments and red are the best global fits (1:1 Langmuir binding model) to the data used to calculate the association rate constants and dissociation rate constants.

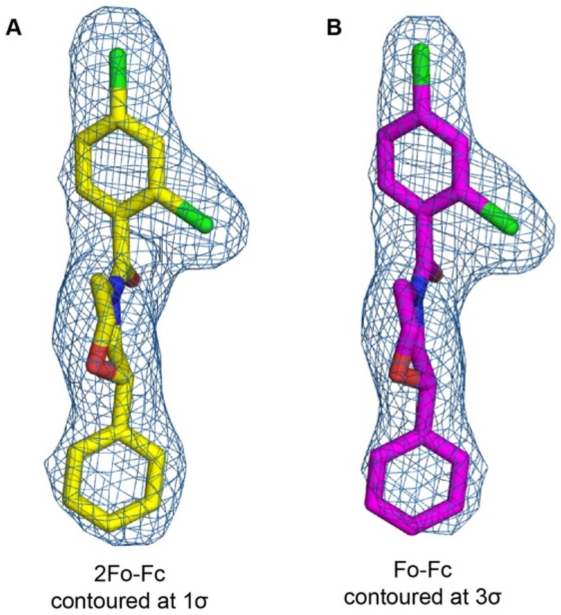


**Fig. S6.** Electron density maps for F0045(S) in complex with H1 PR8 HA. (A) $2F_o$-$F_c$ map (sky blue mesh) generated from the final refined structure of F0045(S) in complex with H1 PR8 HA, contoured at 1σ. Carbon, oxygen, nitrogen, and chlorine are represented in yellow, red, blue, and green, respectively. (B) An unbiased $F_o$-$F_c$ electron density map (sky blue mesh) generated after molecular replacement (MR) using only an apo structure of H1 HA as the MR model, contoured at 3σ. The final refined F0045(S) model is shown superimposed on the unbiased difference map.

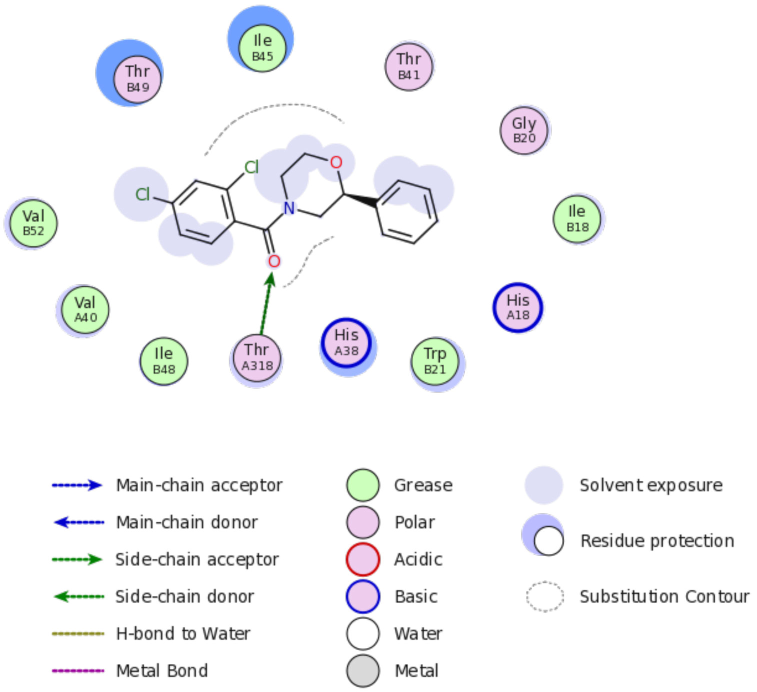


**Fig. S7.** Two-dimensional depiction of the F0045(S) binding sites on influenza H1/PR8 HA. Interactions of F0045(S) with binding-site residues. The arrow indicates a hydrogen bond interaction.

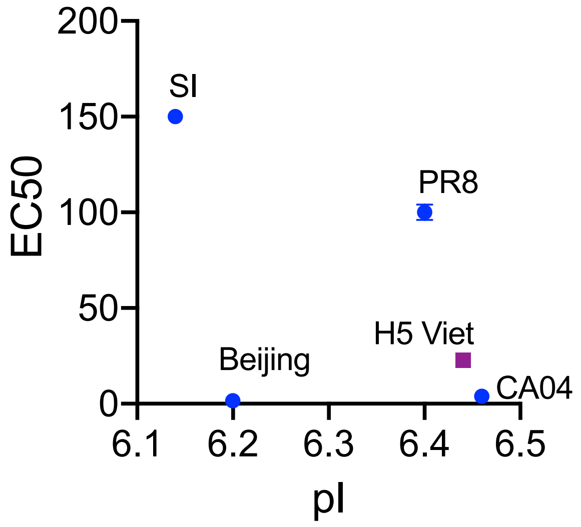


**Fig. S8**. Theoretical pI values of influenza group 1 H1 (blue circle) and H5 (purple square) HAs vs. cellular $EC_{50}$ values.

A

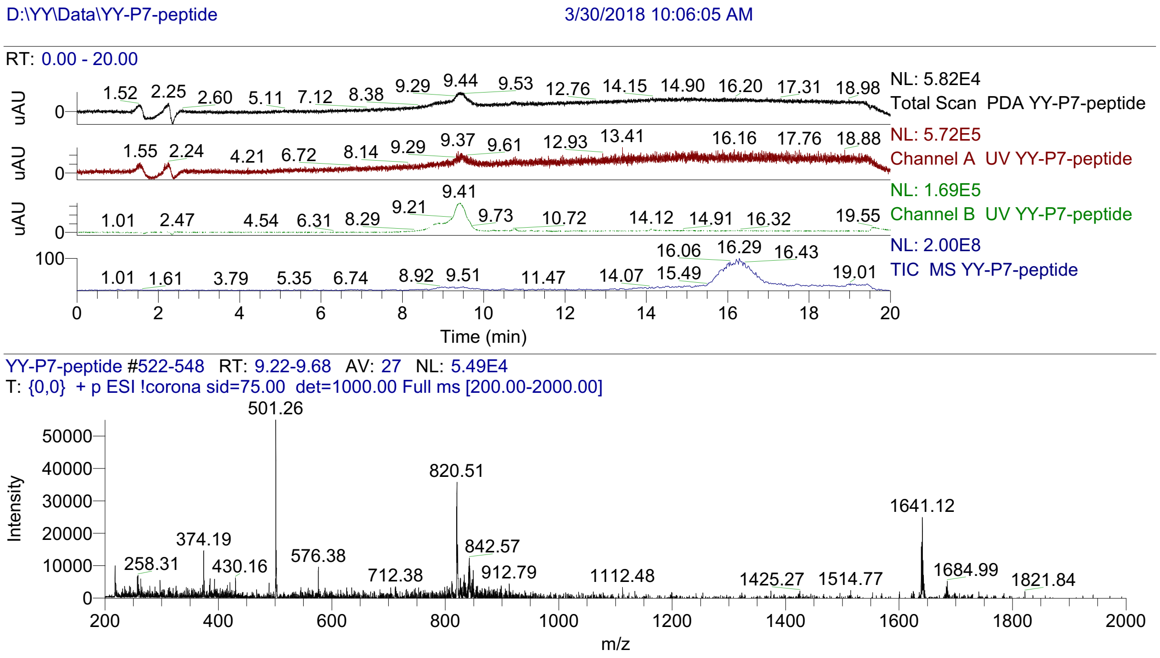


B

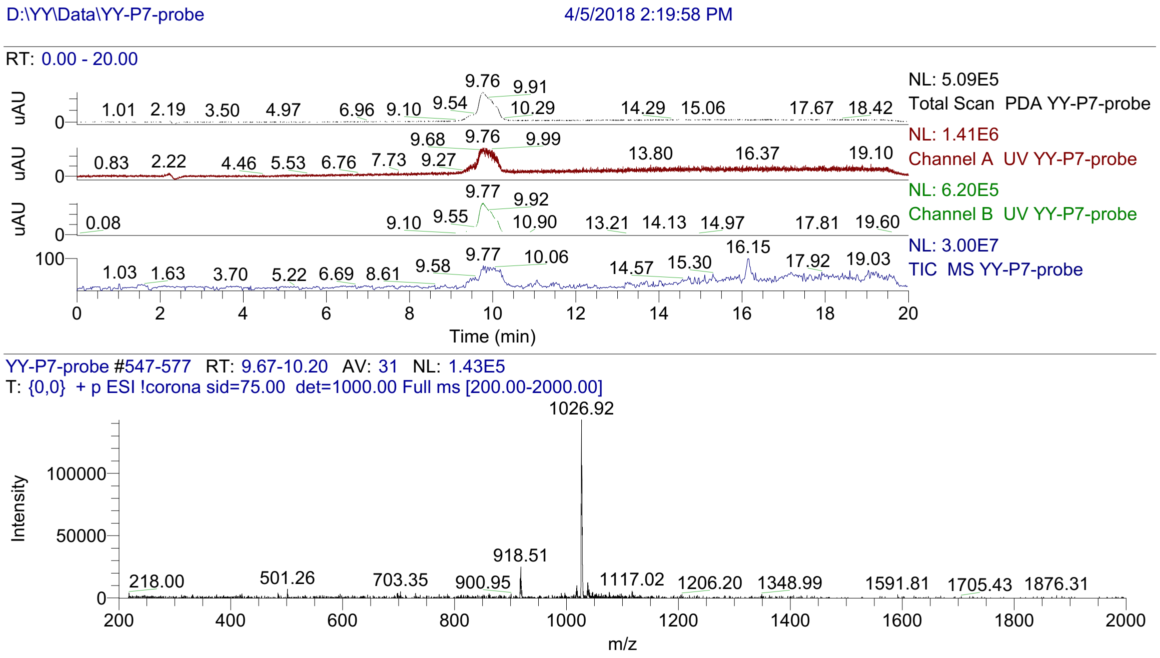


**Fig. S9**. LC-MS characterization of P7 peptide (A) and P7-TAMRA probe (B).

A

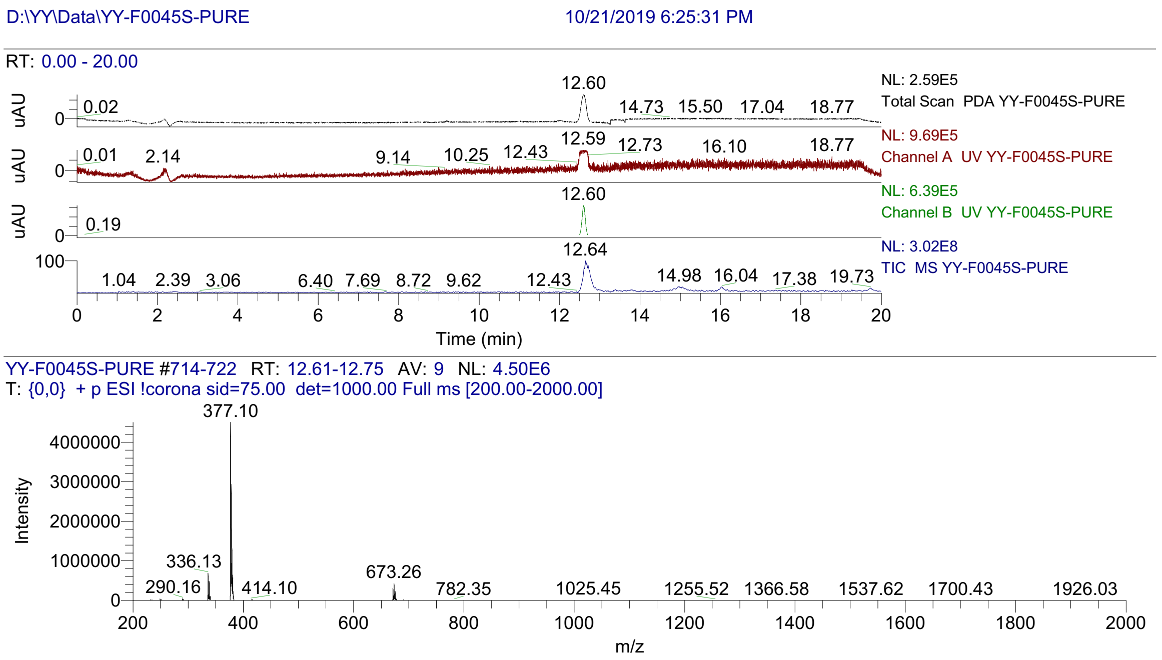


B

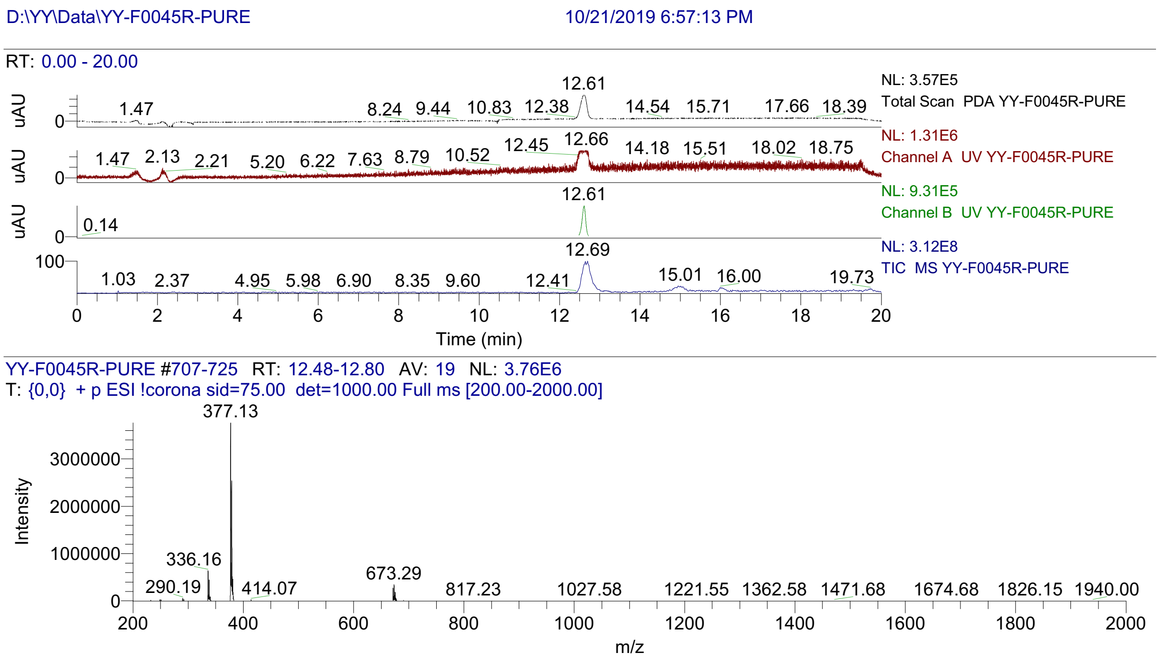


**Fig. S10**. LC-MS characterization of F0045(S) (A) and F0045(R) (B).

A

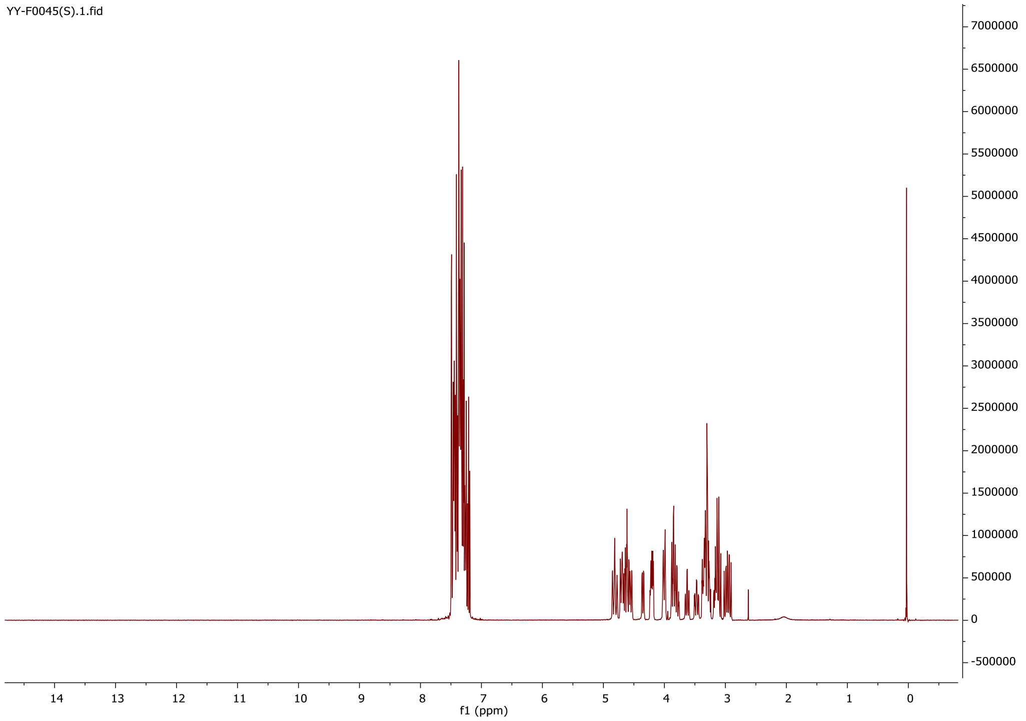


B

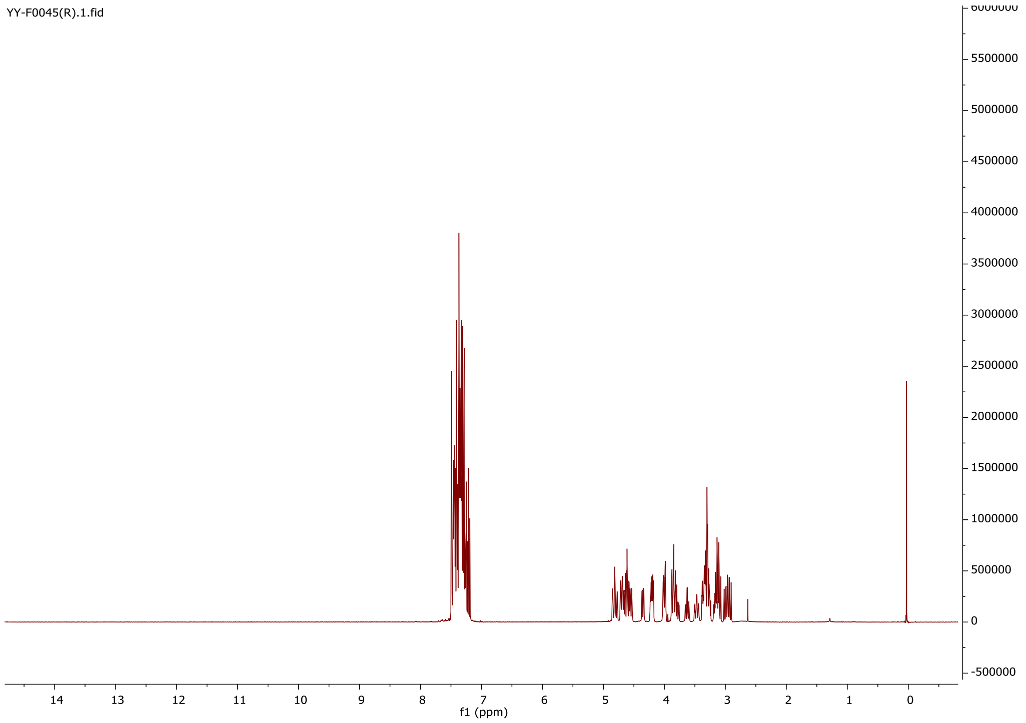


**Fig. S11**. NMR characterization of F0045(S) (A) and F0045(R) (B).

**Table S1**. Data collection and refinement statistics for F0045(S) complex with H1/PR8 HA.

| **X-ray data** | **F0045(S)-H1/PR8** |
|---|---|
| **Data collection** | |
| Beamline | APS-23-IDB |
| Wavelength (Å) | 1.03316 |
| Space Group | $I2_13$ |
| Unit cell (Å,°) | a=b=c=106.1 |
| Resolution range (Å)[a] | 50-2.69 (2.74-2.69) |
| Observations | 381,959 |
| Unique reflections | 19,238 (1923) |
| Completeness (%) | 100(100) |
| I/σ(I) | 27.8 (1.5) |
| $R_{sym}$[b] | 0.15 (1.27) |
| $R_{pim}$[c] | 0.03 (0.37) |
| $CC_{1/2}$[d] | 0.95 (0.67) |
| Redundancy | 19.9 (12.6) |
| **Refinement** | |
| Resolution (Å) | 40.02-2.69 |
| No. reflections[e] | 19,207(1923) |
| $R_{cryst}$[f]/$R_{free}$[g] | 0.22/0.24 |
| No. atoms | |
| Protein | 3922 |
| Ligand/Carbohydrate | 22/95 |
| Water | 36 |
| Wilson B (Å$^2$) | 62 |
| Average B value (Å$^2$) | |
| Protein | 65 |
| Ligand | 44 |
| Water | 56 |
| RMSD from ideal geometry | |
| Bond length (Å) | 0.002 |
| Bond angle (º) | 0.4 |
| Ramachandran Statistics (%)[h] | |
| Favored | 96.5 |
| Outliers | 0.2 |
| PDB ID | 6WCR |

[a]Parentheses refer to outer shell statistics.

[b]$R_{sym} = \Sigma_{hkl} \Sigma_i | I_{hkl,i} - \langle I_{hkl}\rangle | / \Sigma_{hkl} \Sigma_i I_{hkl,i}$, where $I_{hkl,i}$ is the scaled intensity of the $i^{th}$ measurement of reflection *h, k, l*, and $\langle I_{hkl}\rangle$ is the average intensity for that reflection.

[c]$R_{pim} = \Sigma_{hkl} (1/(n-1))^{1/2} \Sigma_i | I_{hkl,i} - \langle I_{hkl}\rangle | / \Sigma_{hkl} \Sigma_i I_{hkl,i}$, where n is the redundancy.

[d]$CC_{1/2}$ = Pearson Correlation Coefficient between two random half datasets.

[e] Value in parentheses refer to number of reflections in test set.

[f]$R_{cryst} = \Sigma_{hkl} | F_o - F_c | / \Sigma_{hkl} | F_o |$ x 100, where $F_o$ and $F_c$ are the observed and calculated structures factors.

[g]$R_{free}$ was calculated as for $R_{cryst}$, but on a test set of 5% of the data excluded from refinement.

[h]Calculated using MolProbity.
